# Genotoxicity and 90-day oral toxicity of a monk fruit (*Siraitia grosvenorii*) mogroside preparation (≥95% mogrosides) produced by microbial fermentation

**DOI:** 10.64898/2026.09.04.749478

**Authors:** Christine Nicole S. Santos, S M Sulaiman, Sanjaykumar M. Paneliya, Martyna Pawluk, A Srikanth, Venkata Sai Prakash Chaturvedula

**Affiliations:** Manus Bio Inc., 43 Foundry Ave., Ste. 230, Waltham, MA 02453, United States; Adgyl Lifesciences Private Limited, 21 & 22, Phase-II, Peenya Industrial Area, Bengaluru 560 058, Karnataka, India

**Keywords:** **Monk Fruit Sweeteners**, Mogrosides, Mogroside V, *Siraitia grosvenorii*, Subchronic toxicity, NOAEL

## Abstract

Here we report genotoxicity and 90-day oral toxicity evaluations of a high-purity, fermentation-derived mogroside preparation rich in mogroside V, the principal sweetener of monk fruit (*Siraitia grosvenorii*). The test article (≥95% total mogrosides, 70.3% mogroside V) is produced by a modified *Escherichia coli* from glucose, offering a higher-purity alternative to traditional monk fruit extracts. The test article was non-mutagenic in a bacterial reverse mutation test (OECD TG 471) and non-clastogenic in an in vitro human lymphocyte micronucleus test (OECD TG 487), up to the maximum recommended concentration. In the 90-day study (OECD TG 408), the test article was given by daily oral gavage at 0, 500, 1000, and 2000 mg/kg body weight/day to Sprague-Dawley rats. There were no deaths and no test article-related effects on clinical signs, ophthalmology, functional observational battery, body weight, food consumption, clinical pathology, thyroid hormones, oestrous cyclicity, or sperm parameters. Minor liver-weight increases lacking any clinical chemistry or histopathological correlates were determined to be non-adverse. No effects were seen on testis weight, sperm endpoints, or spermatogenesis. All other statistically significant differences were minor and considered incidental. The no-observed-adverse-effect level (NOAEL) was 2000 mg/kg body weight/day, the highest dose tested, supporting its safety as a food ingredient.

**Highlights:**

- First safety data on a fermentation-derived mogroside V-rich sweetener.
- The test article was non-mutagenic (Ames) and non-clastogenic (in vitro micronucleus).
- No adverse effects were found in rats at levels up to 2000 mg/kg bw/day for 90 days.
- Importantly, no effects were observed on male fertility end-points.
- The no-observed-adverse-effect level (NOAEL) was 2000 mg/kg bw/day, the highest dose tested.

## 1. Introduction

Excess dietary sugar is a well-established contributor to obesity, type 2 diabetes, and cardiovascular disease, and reducing added sugar in foods and beverages has become a central public-health and reformulation objective, reinforced by front-of-package labelling and sugar-reduction policies across many jurisdictions. Non-caloric high-potency sweeteners are a principal tool for meeting this objective, and demand has shifted increasingly toward nature-derived options. Mogrosides, the cucurbitane-type triterpene glycosides responsible for the intense sweetness of the fruit of *Siraitia grosvenorii* (monk fruit; luo han guo), are among the group of natural high-potency sweeteners of commercial interest: they activate the sweet-taste receptor without being metabolized as carbohydrate, deliver sweetness with no glycemic response, and are heat- and pH-stable, making them suitable across beverages, dairy, and baked applications. All mogrosides share a common cucurbitane aglycone, mogrol, and differ in the number and linkage of attached glucose units. These structural differences influence sweetness intensity, sensory profile, solubility, and metabolic fate. Mogroside V is the most abundant mogroside in the fruit and the principal sweet constituent of commercial monk fruit extracts (Chen et al., 2024; Itkin et al., 2016; Matsumoto et al., 1990).

Monk fruit extracts and juice concentrates have a long history of dietary use in Asia and are Generally Recognized as Safe (GRAS) for use as sweeteners in the United States, supported by rodent and canine toxicity data (Marone et al., 2008; Qin et al., 2006; Jin et al., 2007). However, realizing this potential at the scale that sugar reduction demands has been limited by sweetener supply. The mogroside content of the fruit is low (approximately 0.5-0.65% mogroside V and 3-3.8% total mogrosides per unit dry fruit), the crop is harvested once per year and hand-picked within narrow ripeness windows, and cultivation is concentrated almost entirely in Guangxi province in southern China, which depends on a specific subtropical microclimate that has proven difficult to reproduce elsewhere (Chen et al., 2024; EFSA, 2019). The resulting cost and single-region supply concentration constrain how broadly monk fruit can be deployed for reformulation and expose the supply chain to weather, agricultural, and geopolitical disruption. Fermentation addresses these constraints by reconstructing the mogroside biosynthetic pathway in a microbial host, so that inexpensive carbohydrate feedstock is converted directly into the target sweeteners in fermentors, decoupled from climate, arable land, exposure to toxic pesticides and regional crop limitations (Chen et al., 2024; Itkin et al., 2016). The preparation evaluated here is manufactured by fermentation using a modified strain of *E. coli* K-12 with glucose as the carbon source, followed by downstream purification, yielding a defined ingredient of at least 95% total mogrosides. This route offers two advantages relevant to safety as well as supply. First, it provides a supply chain independent of agricultural monk fruit sourcing, enabling scalable and geographically resilient production. Second, and importantly for hazard characterization, it yields a highly consistent mogroside profile and defined composition than typical monk fruit extracts, which are heterogeneous mixtures that co-extract non-mogroside plant constituents and vary in composition with cultivar, harvest, and growing conditions. Because the fermentation-produced mogrosides are structurally identical to those occurring in *S. grosvenorii*, the existing safety knowledge for these constituents remains relevant for safety assessment, while the higher purity reduces the co-extracted material whose presence can complicate the interpretation of toxicological findings.

To support the safety assessment of this high-purity mogroside preparation intended for use as a sweetening ingredient, a battery of GLP-compliant studies was conducted: a bacterial reverse mutation (Ames) test, an in vitro mammalian cell micronucleus test, and a 90-day repeated-dose oral toxicity study in Sprague-Dawley rats, performed in accordance with OECD Test Guidelines 471, 487, and 408, respectively. Dose levels were selected on the basis of a preceding 14-day dose range-finding study in which no adverse effects were observed at doses up to 3000 mg/kg body weight/day. Unlike the dietary designs used in most prior mogroside studies, the test article was administered by oral gavage, delivering a defined daily dose without diluting the caloric density of the diet, thereby minimizing potential palatability-related and caloric-dilution effects that may complicate interpretation of body-weight changes in dietary sweetener studies. All four studies were conducted under GLP at Adgyl Lifesciences Private Limited (Bengaluru, India), an AAALAC-accredited test facility.

## 2. Materials and methods

### 2.1. Test article

The test article was a high-purity mogroside preparation (Lot No. MAMV2506261, Manus Bio, Inc.) manufactured to a specification of ≥95% total mogrosides. In the lot used for all studies reported here, total mogrosides were 96.5%, of which mogroside V (CAS No. 88901-36-4) was predominant at 70.3%. The remainder comprised mogrosides bearing four or more glucose units, including siamenoside I and mogroside VI, at 17.9%, and minor mogrosides bearing three or fewer glucose units, including mogroside III, at 8.3%; constituents are grouped by degree of glycosylation because all mogrosides are deglycosylated to the common aglycone mogrol before absorption. Moisture, determined as loss on drying, was 4.7%, and ash, residual protein and residual host-cell DNA were each below the limit of quantitation. The test article was manufactured by fermentation using a modified strain of *E. coli* K-12 with glucose as the carbon source, followed by downstream recovery and purification. The mogrosides present in the preparation were determined to be structurally identical to those naturally present in *S. grosvenorii* (monk fruit). The mogroside preparation was stored at the test facility at ambient temperature (+15 to +25 °C), protected from light. The identity and purity of the test article were established by Manus Bio using validated methodologies and were not independently verified at the test facility.

### 2.2. Bacterial reverse mutation (Ames) test

Mutagenic potential was assessed under GLP compliance in accordance with OECD Test Guideline 471 using Salmonella typhimurium strains TA98, TA100, TA1535, and TA1537 and *E. coli* WP2uvrA (pKM101), with and without an exogenous metabolic activation system (S9 fraction prepared from the livers of rats induced with sodium phenobarbitone and beta-naphthoflavone). The test article was dissolved in sterile water as the vehicle. A preliminary toxicity test was conducted over a wide concentration range, after which the test article was evaluated in triplicate at 50, 158, 500, 1581, and 5000 µg/plate. An initial assay used the direct plate-incorporation method and a confirmatory assay used the pre-incubation method. Concurrent vehicle (sterile water) and strain-specific positive controls (2-aminoanthracene with S9; 2-nitrofluorene, sodium azide, 9-aminoacridine, and 4-nitroquinoline-1-oxide without S9) were included. Revertant colonies were counted and the mean and standard deviation calculated for each concentration and control. A test article was considered mutagenic if it produced a concentration-related, reproducible increase in revertants of at least 2-fold (TA98, TA100, WP2uvrA) or 3-fold (TA1535, TA1537) over the concurrent vehicle control. No formal statistical analysis was applied; evaluation was based on these fold-increase criteria together with comparison of revertant counts against the laboratory historical vehicle-control range, in accordance with OECD Test Guideline 471.

### 2.3. In vitro mammalian cell micronucleus test

Clastogenic and aneugenic potential was assessed under GLP compliance in accordance with OECD Test Guideline 487 in cultured primary human peripheral blood lymphocytes obtained from healthy, non-smoking female donors and stimulated with phytohaemagglutinin approximately 48 h before treatment. The test article was dissolved in ultrapure water. Following a preliminary cytotoxicity test, the definitive assay comprised three independent experiments: a 3-h treatment with metabolic activation (+S9), a 3-h treatment without activation (-S9), and a 24-h treatment without activation (-S9), each at duplicate concentrations of 1250, 2500, and 5000 µg/mL (the top concentration corresponding to the OECD/ICH limit applied). Cytokinesis was blocked with cytochalasin B, and cultures were harvested approximately 24 h after the start of treatment. Cytotoxicity was determined from the cytokinesis-block proliferation index (CBPI). Micronuclei were scored in 2000 binucleated cells per concentration (1000 per replicate), and micronucleus frequencies in treated cultures were compared with the concurrent vehicle control. Concurrent positive controls were cyclophosphamide (+S9) and mitomycin C and colchicine (-S9).

### 2.4. Fourteen-Day Dose Range-Finding Study

A 14-day repeated-dose oral gavage study (Study No. AD-N1212) was conducted with reference to OECD Test Guideline 407 in order to establish dose levels for the definitive study; because the study was a pre-study to OECD Test Guideline 408, the duration of treatment and the pathology parameters followed the approved study plan rather than the 28-day guideline. Four groups of five male and five female Sprague-Dawley rats received the test article in ultrapure water by once-daily oral gavage at 0 (vehicle control), 750, 1500, or 3000 mg/kg body weight/day, at a dose volume of 10 mL/kg. Formulations prepared for Day 1 were analyzed for test article concentration and all were within the acceptance limits. Rats were observed for mortality, morbidity, and clinical signs throughout the treatment period, and body weight and food consumption were recorded on Days 1, 4, 8, 11, and 14. On Day 15, following overnight fasting with water available, all rats were subjected to clinical pathology investigations (haematology, coagulation, and clinical chemistry) and a detailed necropsy, and the specified organs were weighed and preserved. Histopathological examination was not performed, at the discretion of the study pathologist and the sponsor, and the preserved tissues were discarded before the report was finalized. Continuous data were analyzed as described in Section 2.5.8. Animal welfare, ethical approval, and reporting statements applicable to in-life studies are given in Section 2.5.1.

### 2.5. 90-Day Oral Toxicity Study

#### 2.5.1. Animals and husbandry

Male and female Sprague-Dawley rats [*Rattus norvegicus*] were obtained from Hylasco Biotechnology (India) Pvt. Ltd. (Telangana, India). Rats were 7-8 weeks old at the start of treatment, with group mean body weights of 256.9-262.4 g (males) and 201.1-204.1 g (females); individual body weights were within ±20% of the sex mean. Females were nulliparous and non-pregnant. After a 5-day acclimatization period, animals were randomized to groups by body weight stratification (Provantis, Instem LSS). Rats were housed two per cage by sex in solid-floor polysulfone cages with corn-cob bedding and polycarbonate enrichment huts, under controlled conditions (19.6-24.9 °C; relative humidity 52-65%; 12 h light/12 h dark). A pelleted maintenance diet (Altromin) and filtered, UV-treated water were provided ad libitum. Both in-life studies were conducted at Adgyl Lifesciences Private Limited (Bengaluru, India), a test facility accredited by AAALAC International. Adgyl’s accreditation is assessed against the Guide for the Care and Use of Laboratory Animals (National Research Council, 2011), and both studies were conducted within the scope of that accredited program and in accordance with the Guide. All procedures were additionally conducted in accordance with the guidelines of the Committee for the Control and Supervision of Experiments on Animals (CCSEA), India, and each study protocol was reviewed and approved by the test facility Institutional Animal Ethics Committee, operating under CCSEA oversight, prior to initiation: Approval No. 209/Aug-2025 for the 14-day dose range-finding study and Approval No. 210/Aug-2025 for the 90-day study, both dated 14 August 2025. Both studies are reported in accordance with the ARRIVE 2.0 guidelines..

#### 2.5.2. Study design and dose administration

The study was conducted under GLP compliance in accordance with OECD Test Guideline 408 (Repeated Dose 90-Day Oral Toxicity Study in Rodents). Four groups of 10 males and 10 females each received the test article by once-daily oral gavage at 0 (vehicle control, G1), 500 (G2), 1000 (G3), or 2000 (G4) mg/kg body weight/day for 90 consecutive days. Dose levels were based on the results of the 14-day range-finding study. The vehicle was ultrapure water, selected because the test article formed a uniform dispersion in it; homogeneity and stability of the formulations were established at 1.00 and 500.10 mg/mL (stable for 7 days at room temperature and 18 days refrigerated). Formulations were administered at a constant volume of 10 mL/kg, with concentrations of 50, 100, and 200 mg/mL for the low-, mid-, and high-dose groups, respectively, and were kept homogeneous by continuous magnetic stirring during dosing. Dose volumes were adjusted to the most recent body weight. Group sizes were those specified by OECD test guidelines and the approved study plans, which prescribe group sizes sufficient to detect toxicologically meaningful effects in a repeated-dose screening design. The experimental unit was the individual animal, the test article being administered individually by gavage, except food consumption, which was measured cage-wise and expressed as g/rat/day.

#### 2.5.3. Dose formulation analysis

Formulations prepared on Day 1 and once each during Months 2 and 3 were sampled for concentration verification using an HPLC method validated under a separate study. Acceptance criteria were a mean measured concentration within ±15% of nominal and a relative standard deviation (RSD) <10%. All analysed samples met these criteria; overall mean recoveries ranged from 94.67% to 105.02% with %RSD <4.0%, confirming accurate and homogeneous dosing.

#### 2.5.4. In-life observations

All rats were observed twice daily for morbidity and mortality. General clinical signs were recorded, and detailed clinical examinations (skin, fur, eyes, mucous membranes, secretions/excretions, autonomic activity, gait, posture, handling response, and abnormal behaviour) were performed prior to dosing on Day 1 and weekly thereafter. Ophthalmological examinations (direct ophthalmoscopy following tropicamide-induced mydriasis) were conducted pretest and at the end of treatment (Day 88). Body weights and food consumption were recorded weekly. Vaginal smears were examined prior to necropsy to record the stage of the oestrous cycle in females. No animals were excluded from either study and no data were excluded from analysis. Assessors were not blinded to treatment group; in-life observations, clinical pathology, and histopathological evaluation were performed with knowledge of dose group, in accordance with standard practice for GLP repeated-dose toxicity studies.

#### 2.5.5. Functional observational battery

A functional observational battery (FOB) was performed during Week 13 (Day 86 for males; Day 88 for females), comprising home-cage observations, handling observations, open-field observations, sensory reactivity, landing hindlimb foot splay, fore-and hindlimb grip strength, body temperature, and motor (locomotor) activity.

#### 2.5.6. Clinical pathology

Blood and urine were collected at termination (Day 91) after overnight fasting. Haematology (ADVIA 2120i) included red blood cell count, haemoglobin, haematocrit, mean cell volume (MCV), mean cell haemoglobin (MCH), mean cell haemoglobin concentration (MCHC), reticulocytes, total and differential leukocyte counts, platelets, mean platelet volume, and red cell morphology indices. Coagulation parameters (prothrombin time, PT; activated partial thromboplastin time, APTT) were measured on citrated plasma (Ceveron Alpha). Clinical chemistry (Beckman Coulter AU480) included alanine aminotransferase (ALT), aspartate aminotransferase (AST), alkaline phosphatase (ALP), gamma-glutamyl transferase (GGT), creatine kinase (CK), glucose, blood urea nitrogen, creatinine, total bilirubin, total protein, albumin, globulin (calculated), albumin:globulin (A/G) ratio, total, high-density lipoprotein (HDL) and low-density lipoprotein (LDL) cholesterol, triglycerides, inorganic phosphorus, calcium, sodium, potassium, and chloride. Serum total triiodothyronine (T3), thyroxine (T4), and thyroid-stimulating hormone (TSH) were determined by immunoassay (Calbiotech and Bioelsa kits). Urine was collected overnight (Day 90-91) and evaluated for specific gravity, pH, nitrite, protein, glucose, ketones, urobilinogen, bilirubin, erythrocytes, leukocytes, volume, colour, clarity, and microscopic sediment.

#### 2.5.7. Necropsy, organ weights, sperm analysis, and histopathology

All rats were fasted overnight, weighed (terminal fasting body weight), euthanized under isoflurane, exsanguinated, and subjected to a full macroscopic examination. A standard set of organs was weighed, and organ-to-terminal-body-weight and organ-to-brain-weight ratios were calculated; paired organs were weighed together. A comprehensive list of tissues was preserved in 10% neutral-buffered formalin (testes and eyes in Bouin’s/appropriate fixative). Sperm motility was assessed in all males by computer-assisted sperm analysis (Hamilton-Thorne). Sperm morphology, cauda epididymal sperm counts, and detergent- and homogenization-resistant testicular spermatid counts were determined in the vehicle control (G1) and high-dose (G4) groups only, matching the histopathological sampling strategy described below. Histopathological examination was performed on the full tissue list from the vehicle control (G1) and high-dose (G4) groups; in the absence of test article-related findings in G4, lower-dose tissues were not examined. Testicular staging of spermatogenesis was included.

#### 2.5.8. Statistical analysis

Continuous data (body weight, body weight change, body temperature, grip strength, food consumption, organ weights and ratios, haematology, coagulation, clinical chemistry, foot splay, and motor activity) were assessed for homogeneity of variance (Levene test) and normality (Shapiro-Wilk). Homogeneous, normally distributed data were analysed by one-way analysis of variance (ANOVA); non-homogeneous or non-normal data were transformed prior to ANOVA. Where ANOVA was significant, treated groups were compared with control by Dunnett’s test. All hypothesis testing was two-sided at the 5% significance level (p < 0.05).

## 3. Results

### 3.1. Bacterial reverse mutation (Ames) test

The test article was soluble in sterile water and produced no precipitation on the plates up to 5000 µg/plate. In the preliminary toxicity test, no reduction in the bacterial background lawn or in revertant numbers was observed at any concentration, so the definitive assays were conducted up to the maximum recommended concentration of 5000 µg/plate. In both the initial (plate-incorporation) and confirmatory (pre-incubation) assays, the test article produced no concentration-related, reproducible increase in revertant colony counts in any of the five tester strains, either with or without metabolic activation; mean revertant counts at all concentrations were comparable to the concurrent vehicle control and within the laboratory historical control ranges (Table 1). The strain-specific positive controls each produced the expected large increase in revertants (greater than 3-fold), confirming assay validity and the activity of the S9 mix. Under the conditions of this study, ≥95% mogrosides were concluded to be non-mutagenic in the bacterial reverse mutation assay up to the highest OECD TG 471–recommended dose of 5,000 µg/plate.

**Table 1.**
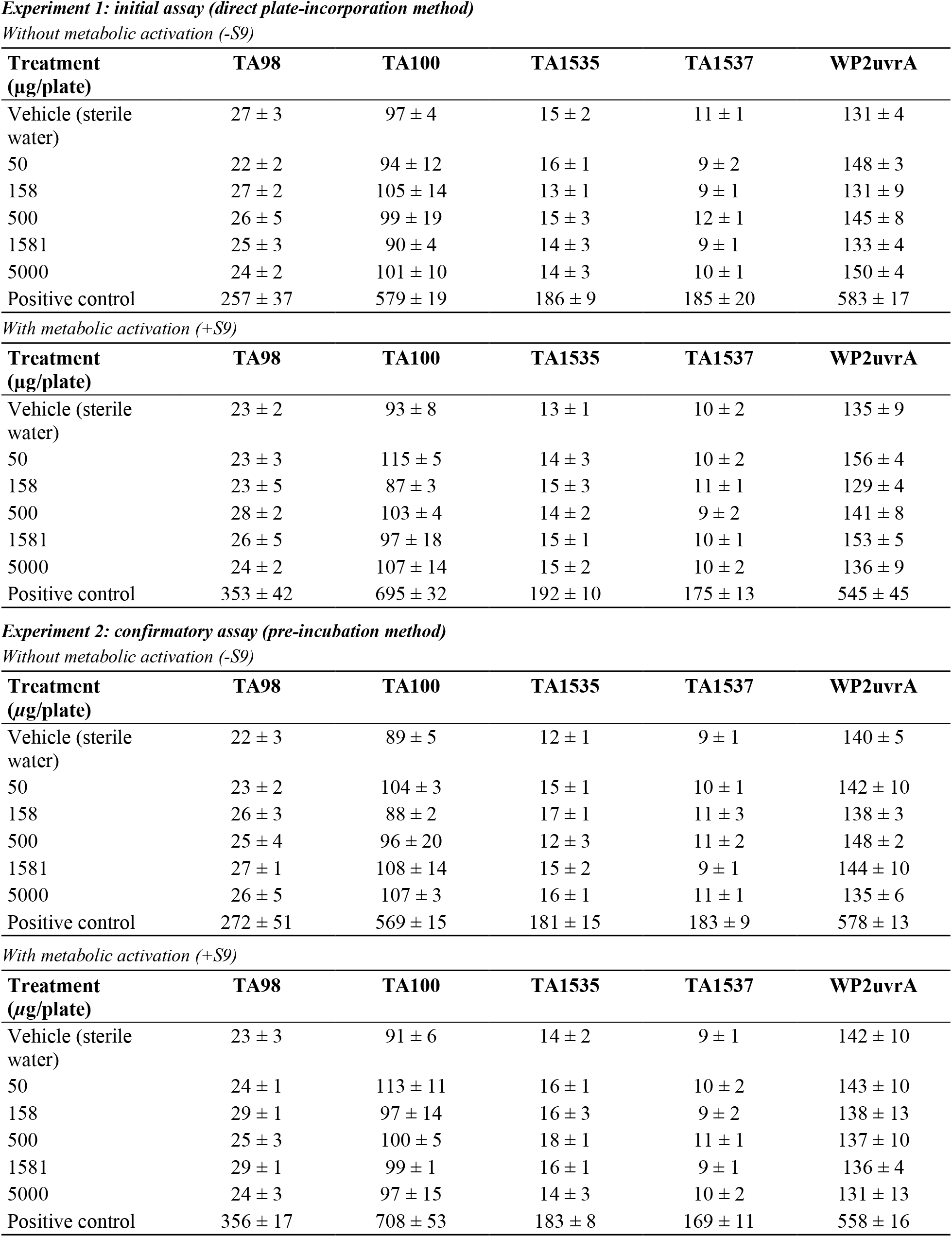
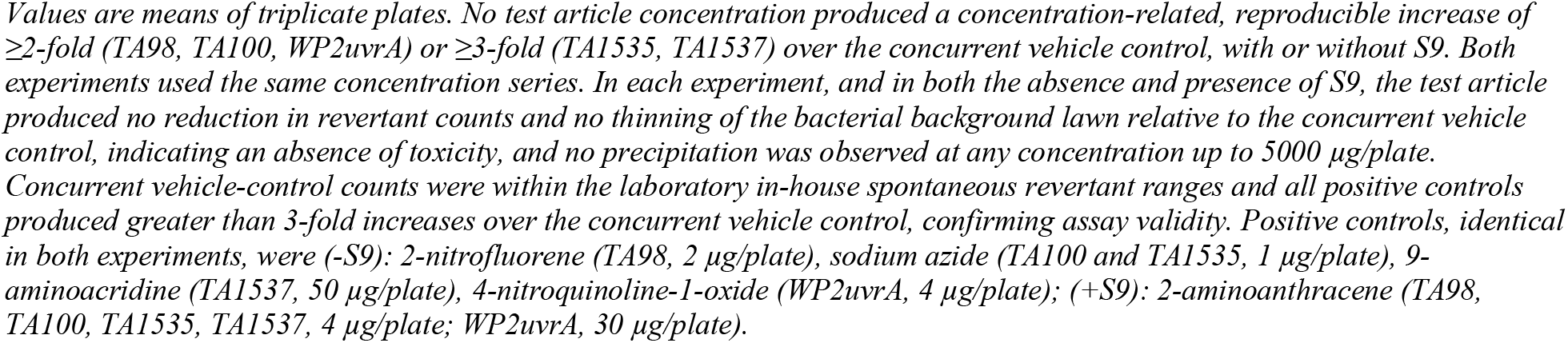
Bacterial reverse mutation (Ames) test: mean revertant colonies per plate (mean ± SD of triplicate plates) in the initial (plate-incorporation) and confirmatory (pre-incubation) assays.

### 3.2. In vitro mammalian cell micronucleus test

The test article did not appreciably alter the pH or osmolality of the treatment medium at any concentration. In the definitive assay, graded cytotoxicity was achieved at the top concentration (5000 *µ*g/mL), with reductions in CBPI corresponding to 38%, 44%, and 46% cytotoxicity in the 3-h +S9, 3-h -S9, and 24-h - S9 experiments, respectively, confirming adequate exposure. Across all three experiments, the frequency of micronucleated binucleated cells in test article-treated cultures was not statistically increased relative to the concurrent vehicle control at any concentration, with or without metabolic activation, and remained within the historical control range (Table 2). The positive controls (cyclophosphamide, mitomycin C, and colchicine) each produced a statistically significant increase in micronucleated cells, confirming the sensitivity of the test system. The test article was therefore non-clastogenic and non-aneugenic under the conditions of the test.

**Table 2.** In vitro micronucleus test in human lymphocytes: frequency of micronucleated binucleated cells (% MNBN) and cytotoxicity (per pooled 2000 cells/concentration).

| Concentration ($\mu\text{g}/\text{mL}$) | 3h +S9 % MNBN | 3h -S9 % MNBN | 24h -S9 % MNBN | Cytotox. at top conc. (%) |
| --- | --- | --- | --- | --- |
| Vehicle (sterile water) | 1.10 | 1.05 | 1.30 | - |
| 1250 | 0.75 | 0.85 | 0.80 | - |
| 2500 | 0.85 | 0.95 | 1.00 | - |
| 5000 | 0.95 | 0.90 | 1.10 | 38 / 44 / 46 |
| Positive control | 10.90* | n/a | 10.85 / 10.60* | - |

\* $p < 0.05$ vs. vehicle control. No test article concentration significantly increased % MNBN relative to the concurrent vehicle control in any experiment; all values were within the laboratory historical vehicle-control ranges (% micronucleated cells; HPBL, 2018-2024, $n = 169$ ): 3h +S9, 0.50-1.30 (mean 0.94); 3h -S9, 0.60-1.45 (mean 0.93); 24h -S9, 0.65-1.40 (mean 0.99). Cytotoxicity (CBPI-based) at the top concentration (5000 $\mu\text{g}/\text{mL}$ ) was 38%, 44%, and 46% in the 3h +S9, 3h -S9, and 24h -S9 experiments, respectively. Positive controls: cyclophosphamide (+S9, 3h); mitomycin C and colchicine (-S9, 24h).

### 3.3. Fourteen-Day Dose Range-Finding Study

There was no mortality and there were no test article-related clinical signs at any dose in either sex. Mean body weights and body weight gains were unaffected in both sexes. Food consumption was unaffected in females; in males at 3000 mg/kg body weight/day a statistically significant reduction of approximately 7-9% was recorded from Days 4 to 14, which was considered to relate to the nature of the test article and to be non-adverse, since the difference was marginal (<10%) and mean body weights were unaffected. No test article-related changes were observed in haematology, coagulation, or clinical chemistry; the statistically significant differences in creatine kinase in males and in total bilirubin in both sexes were considered incidental and of no toxicological relevance. Terminal fasting body weights, organ weights, and organ weight ratios were unaffected in either sex, and the statistically significant increase in adrenal weights in females at 750 and 3000 mg/kg body weight/day were considered incidental in the absence of dose dependency. There were no test article-related macroscopic findings. The study therefore identified no adverse effects up to the highest dose tested, 3000 mg/kg body weight/day, and supported the dose levels selected for the 90-day study.

### 3.4. 90-Day Oral Toxicity Study

#### 3.4.1. Mortality, clinical signs, and ophthalmology

There were no deaths during the study. No test article-related clinical signs were observed at any dose level in either sex, and detailed clinical examinations were unremarkable. Ophthalmological examination revealed no test article-related ocular abnormalities.

#### 3.4.2. Functional observational battery

Home-cage, handling, open-field, and sensory observations, body temperature, and motor activity were comparable to control at all doses. Isolated statistically significant differences in increased forelimb grip strength in males at 500 mg/kg/day, increased hindlimb grip strength in females at 500 mg/kg/day, and decreased hindlimb foot splay in females at 1000 mg/kg/day were small, lacked a dose-response relationship, occurred without corroborative changes in extensor thrust response or muscle tone, and were not considered test article-related.

#### 3.4.3. Body weight and food consumption

Mean body weights, body weight gains, and food consumption were unaffected by treatment in both sexes (Figs. 1 and 2). Scattered statistically significant differences in interval body weight gain (males: increased on Days 15-22 at 2000 mg/kg/day and decreased on Days 43-50 at 500 mg/kg/day; females: increased on Days 22-29 at 500 mg/kg/day) and in interval food consumption in females (7-9% increases at 2000 mg/kg/day on Days 29-36 and at 500 mg/kg/day on Days 36-43) were inconsistent across dose, time, and sex, were not associated with changes in mean body weight, and were considered incidental. Overall (Day 1-90) body weight gain did not differ significantly from control in either sex.

**Fig. 1.**
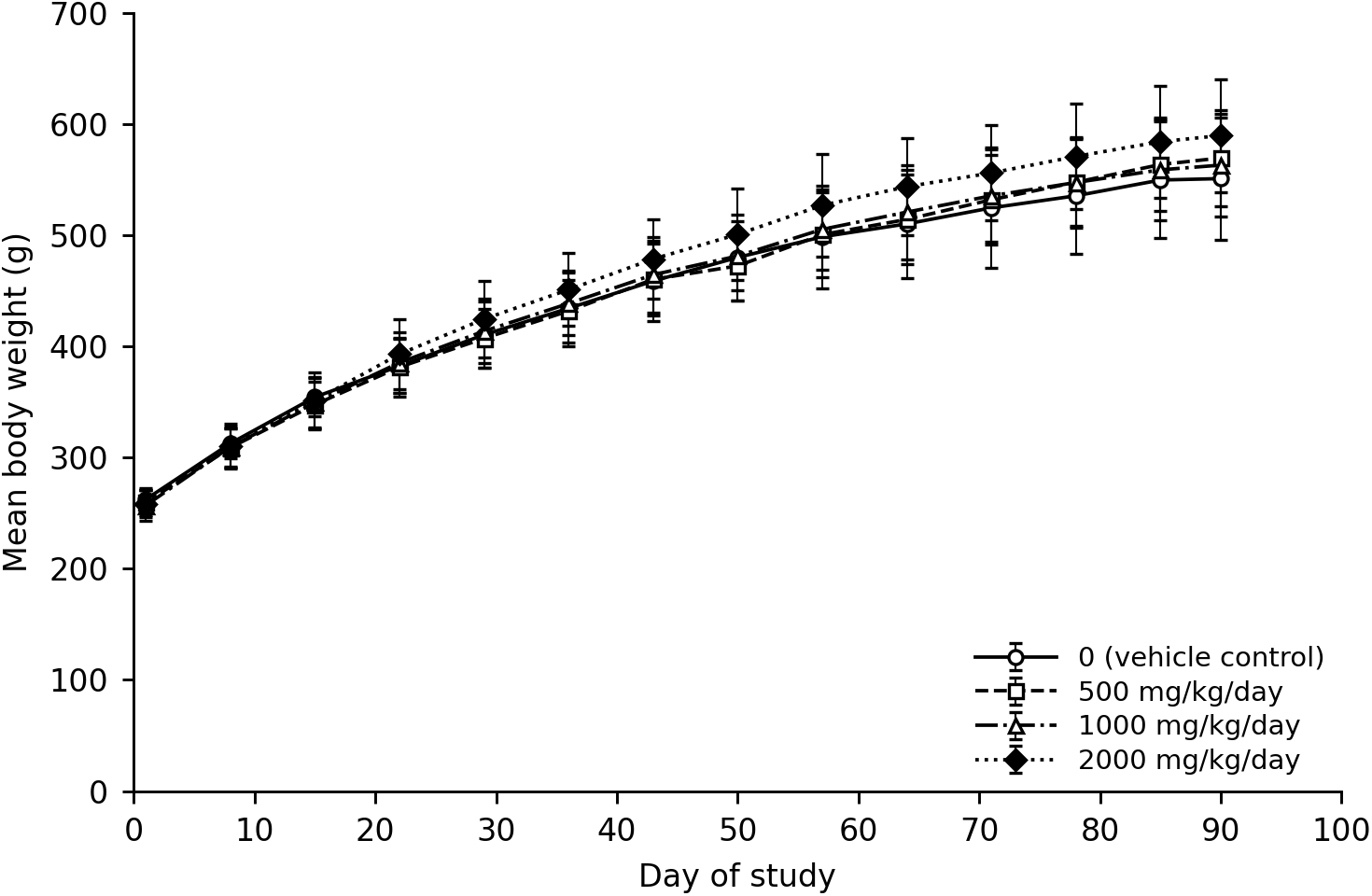
Weekly mean body weights of male rats administered the test article by oral gavage at 0, 500, 1000, or 2000 mg/kg body weight/day for 90 days (n = 10/group). Error bars are ± SD (n = 10/group). No treated group differed significantly from control at any interval.

**Fig. 2.**
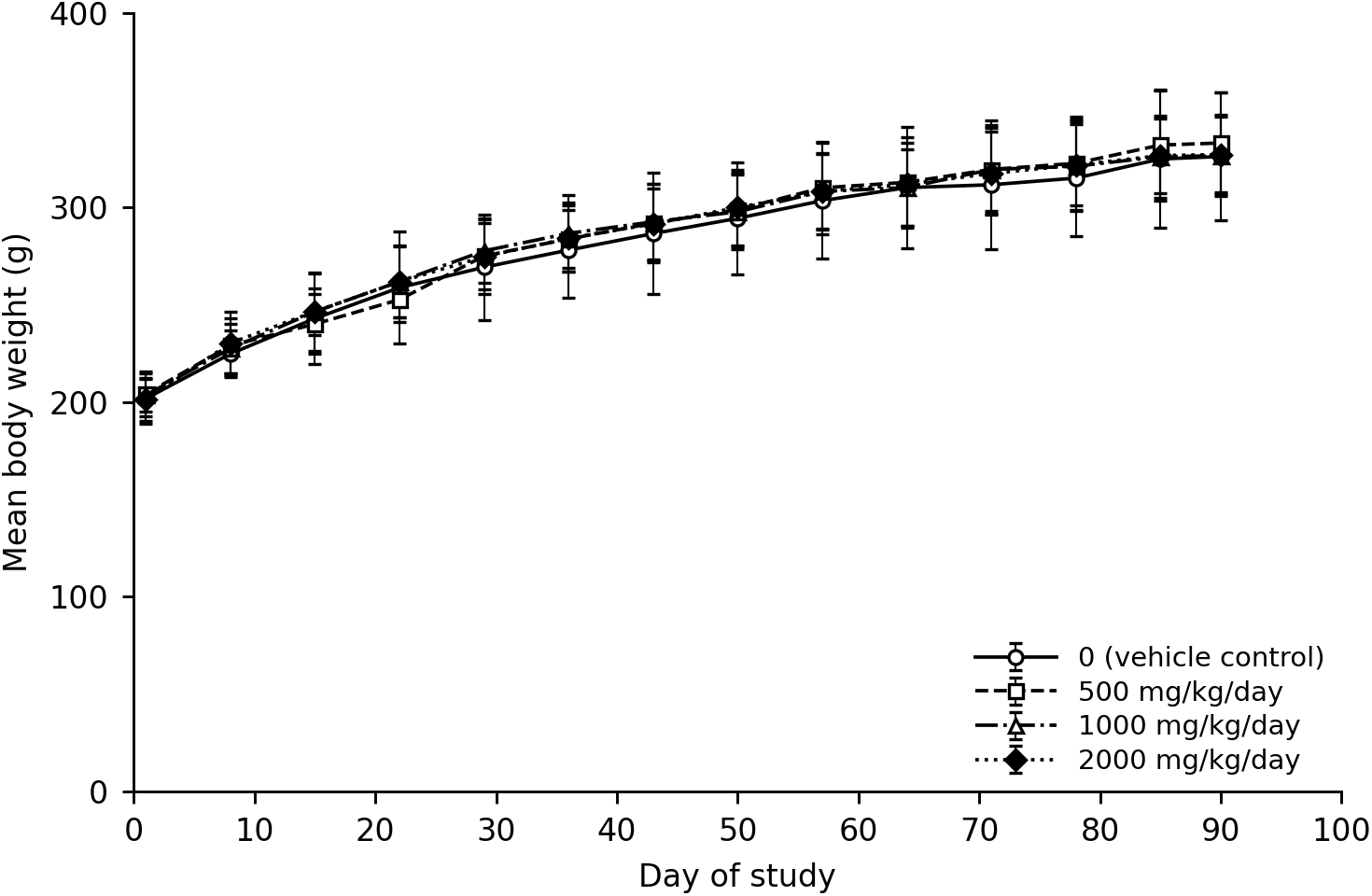
Weekly mean body weights of female rats administered the test article by oral gavage at 0, 500, 1000, or 2000 mg/kg body weight/day for 90 days (n = 10/group). Error bars are ± SD (n = 10/group). No treated group differed significantly from control at any interval.

#### 3.4.4. Oestrous cyclicity

The stage of the oestrous cycle recorded prior to necropsy in treated females was comparable to that of the vehicle control group (data not shown).

#### 3.4.5. Clinical pathology

No test article-related changes were observed in haematology, coagulation, clinical chemistry, thyroid hormones, or urinalysis (Tables 3-5; complete haematology, coagulation, clinical chemistry and urinalysis panels in Supplementary Tables S1a-S1c). In haematology, a modest increase in mean platelet volume was noted in males (7-12%) at all doses and in females (4-5%) at 1000 and 2000 mg/kg/day, without accompanying changes in platelet count; decreased white cell (19%) and lymphocyte (21%) counts occurred in high-dose males; and neutrophil counts were numerically but not statistically significantly higher (28-30%) in mid- and high-dose females. These differences were of small magnitude, lacked a clear dose-response, were within historical control ranges, and had no microscopic correlate at the vehicle control and high-dose groups, the only groups in which histopathology was performed. Tissues from the 500 and 1000 mg/kg/day groups were not examined because no test item-related change was identified at 2000 mg/kg/day. A 22% increase in APTT in high-dose males was likewise isolated, without related changes in PT, platelet count, clinical signs of bleeding, or histopathology. In clinical chemistry, total bilirubin was lower across dose groups in males (11-17%) and females (15-31%) with no correlative findings and values within historical control ranges. A higher T4 in high-dose females occurred without changes in T3 or TSH and without thyroid organ-weight or microscopic correlates. Urine volume was lower at 1000 and 2000 mg/kg/day in males (-39% and -31%) and at 1000 mg/kg/day in females (-44%), with correspondingly higher specific gravity and marginally lower pH at the top dose in both sexes. These differences were not accompanied by any change in blood urea nitrogen, creatinine, or electrolytes, or by any kidney organ-weight or microscopic correlate and are consistent with a difference in urine concentrating status rather than with renal injury. All such findings were considered incidental and not test article-related.

**Table 3.** Selected haematology and coagulation parameters at termination (Day 91; mean ± SD, n = 10/sex/group).

| Parameter (unit) | 0 | 500 | 1000 | 2000 |
| --- | --- | --- | --- | --- |
| RBC ($10^{12}/\text{L}$) – M | 9.08 ± 0.31 | 9.02 ± 0.29 | 8.77 ± 0.32* | 9.11 ± 0.23 |
| HGB (g/L) – M | 168 ± 5 | 168 ± 6 | 168 ± 8 | 172 ± 4 |
| WBC ($10^9/\text{L}$) – M | 10.09 ± 1.48 | 10.20 ± 1.51 | 9.70 ± 1.82 | 8.15 ± 1.89* |
| PLT ($10^9/\text{L}$) – M | 1283 ± 110 | 1295 ± 63 | 1483 ± 304 | 1190 ± 131 |
| MPV (fL) – M | 9.6 ± 0.4 | 10.3 ± 0.5* | 10.5 ± 0.3* | 10.8 ± 0.4* |
| APTT (s) – M | 17.4 ± 2.5 | 17.5 ± 2.6 | 17.4 ± 3.5 | 21.2 ± 2.3* |
| RBC ($10^{12}/\text{L}$) – F | 8.19 ± 0.26 | 8.27 ± 0.38 | 8.25 ± 0.28 | 8.28 ± 0.22 |
| HGB (g/L) – F | 157 ± 5 | 160 ± 4 | 161 ± 6 | 160 ± 4 |
| WBC ($10^9/\text{L}$) – F | 7.06 ± 1.08 | 5.69 ± 1.02* | 5.98 ± 1.07 | 7.62 ± 1.61 |
| MPV (fL) – F | 10.1 ± 0.4 | 10.4 ± 0.3 | 10.6 ± 0.3* | 10.5 ± 0.3* |
| APTT (s) – F | 12.9 ± 1.8 | 11.7 ± 2.4 | 11.0 ± 3.7 | 13.4 ± 1.5 |

**Table 4.** Selected clinical chemistry parameters at termination (Day 91; mean ± SD, n = 10/sex/group).

| Parameter (unit) | 0 | 500 | 1000 | 2000 |
| --- | --- | --- | --- | --- |
| ALT (U/L) – M | 32 ± 5 | 46 ± 38 | 31 ± 5 | 35 ± 11 |
| ALP (U/L) – M | 85 ± 19 | 91 ± 15 | 78 ± 15 | 81 ± 16 |
| Total bilirubin (μmol/L) – M | 2.31 ± 0.36 | 2.05 ± 0.62 | 1.92 ± 0.27 | 1.93 ± 0.45 |
| Total cholesterol (mmol/L) – M | 1.64 ± 0.42 | 1.75 ± 0.28 | 1.52 ± 0.23 | 1.61 ± 0.29 |
| Glucose (mmol/L) – M | 6.38 ± 1.01 | 6.35 ± 0.81 | 6.13 ± 0.44 | 6.66 ± 0.71 |
| Creatinine (μmol/L) – M | 40 ± 2 | 41 ± 4 | 42 ± 4 | 42 ± 4 |
| Total protein (g/L) – M | 71.0 ± 3.1 | 72.6 ± 2.3 | 70.0 ± 2.5 | 72.9 ± 1.7 |
| ALT (U/L) – F | 30 ± 21 | 28 ± 14 | 45 ± 46 | 39 ± 43 |
| ALP (U/L) – F | 57 ± 19 | 39 ± 5 | 36 ± 9* | 43 ± 17 |
| Total bilirubin (μmol/L) – F | 2.96 ± 0.77 | 2.51 ± 0.27 | 2.38 ± 0.40 | 2.04 ± 0.37* |
| Total cholesterol (mmol/L) – F | 2.32 ± 0.47 | 2.44 ± 0.40 | 2.33 ± 0.69 | 2.26 ± 0.47 |
| Creatinine (μmol/L) – F | 45 ± 5 | 48 ± 3 | 50 ± 3* | 46 ± 4 |

**Table 5.** Thyroid hormone concentrations at termination (Day 91; mean ± SD, n = 10/sex/group).

| Parameter (unit) | 0 | 500 | 1000 | 2000 |
| --- | --- | --- | --- | --- |
| T3 (ng/mL) – M | 0.89 ± 0.21 | 0.86 ± 0.19 | 0.67 ± 0.11* | 0.77 ± 0.21 |
| T4 (ng/mL) – M | 73.38 ± 14.75 | 76.75 ± 12.87 | 72.72 ± 11.71 | 69.27 ± 10.87 |
| TSH (ng/mL) – M | 1.12 ± 0.12 | 1.16 ± 0.14 | 1.08 ± 0.16 | 1.13 ± 0.17 |
| T3 (ng/mL) – F | 0.70 ± 0.11 | 0.72 ± 0.18 | 0.75 ± 0.19 | 0.81 ± 0.26 |
| T4 (ng/mL) – F | 46.18 ± 9.07 | 51.51 ± 11.03 | 50.62 ± 10.91 | 61.68 ± 14.94* |
| TSH (ng/mL) – F | 1.02 ± 0.17 | 1.14 ± 0.13 | 1.03 ± 0.19 | 1.05 ± 0.17 |

Dose in mg/kg body weight/day. M, males; F, females. \* $p < 0.05$ vs. control (Dunnett). The lower T3 in males at 1000 mg/kg body weight/day and the higher T4 in females at 2000 mg/kg body weight/day were judged incidental rather than test article-related because neither followed a dose-response relationship, neither was accompanied by a change in any other thyroid hormone in the same sex, and neither had a thyroid organ-weight or histopathological correlate; both values fell within the laboratory historical control ranges.

#### 3.4.6. Organ weights

Terminal fasting body weights were unaffected by treatment in both sexes. Increased liver weights were observed in high-dose males (absolute +13%; organ-to-brain ratio +18%, p < 0.05) and in mid- and high-dose females (absolute +14% and +12%; organ-to-body-weight ratio +13% and +12%, p < 0.05) (Table 6). In the absence of any corresponding clinical chemistry change (e.g., ALT, ALP, GGT, bilirubin) or hepatic histopathological correlate, these liver-weight increases were considered test article-related but non-adverse. Other organ-weight differences, including increased liver weight in low-dose males, increased thymus weight in males, and increased adrenal and ovary weights in females, were minimal and/or lacked a clear dose-response, had no microscopic or clinical pathology correlate, and were within historical control ranges. These findings were considered incidental.

**Table 6.** Terminal fasting body weight and liver weight (mean ± SD, n = 10/sex/group).

| Parameter (unit) | 0 | 500 | 1000 | 2000 |
| --- | --- | --- | --- | --- |
| Terminal fasting bw (g) – M | 514.3 ± 56.4 | 534.6 ± 43.4 | 525.8 ± 41.0 | 553.6 ± 48.8 |
| Liver, absolute (g) – M | 12.11 ± 1.97 | 12.98 ± 1.64 | 12.14 ± 1.27 | 13.64 ± 1.73 |
| Liver/body weight (%) – M | 2.35 ± 0.19 | 2.42 ± 0.14 | 2.31 ± 0.21 | 2.46 ± 0.20 |
| Liver/brain (%) – M | 504.7 ± 95.3 | 551.6 ± 54.9 | 511.9 ± 56.0 | 595.8 ± 78.7* |
| Terminal fasting bw (g) – F | 300.3 ± 33.4 | 310.2 ± 23.6 | 301.9 ± 18.9 | 298.1 ± 17.8 |
| Liver, absolute (g) – F | 7.88 ± 1.01 | 8.40 ± 0.99 | 8.98 ± 0.88 | 8.81 ± 1.07 |
| Liver/body weight (%) – F | 2.62 ± 0.15 | 2.71 ± 0.27 | 2.97 ± 0.18* | 2.95 ± 0.24* |
| Liver/brain (%) – F | 361.1 ± 49.3 | 391.4 ± 52.1 | 405.9 ± 39.8 | 409.3 ± 56.1 |

Dose in mg/kg body weight/day. M, males; F, females. \* $p < 0.05$ vs. control (Dunnett). The liver-weight increases were considered test article-related but non-adverse: they were of small magnitude (12-14% for absolute weight), occurred without any corresponding change in hepatocellular or hepatobiliary clinical chemistry markers (ALT, ALP, GGT, bilirubin), and had no hepatic gross or microscopic correlate.

#### 3.4.7. Sperm analysis

There were no test article-related effects on sperm parameters. The percentages of motile and progressively motile sperm (e.g., progressively motile sperm 64.3-66.8% across groups), testicular spermatid counts, and epididymal sperm counts were comparable between control and treated groups. A higher incidence of abnormal sperm in the high-dose group reflected individual variation (1.0% versus 0.7% in control), was not statistically significant, and had no microscopic correlate in reproductive tissues.

#### 3.4.8. Gross pathology and histopathology

No test article-related gross lesions were observed in either sex. Histopathological examination of control and high-dose tissues revealed no test article-related microscopic findings; because no test article-related change was identified in the high-dose group, tissues from the 500 and 1000 mg/kg/day groups were not examined, as provided for by OECD Test Guideline 408. The few observed changes were consistent with common spontaneous or incidental background findings distributed randomly across groups. Qualitative evaluation of spermatogenic staging was normal, with complete spermatogenic cycles and no evidence of stage-specific arrest.

## 4. Discussion

A bacterial reverse mutation assay, an in vitro mammalian cell micronucleus assay, and a 90-day repeated-dose oral toxicity study were conducted to characterize the safety of a high-purity, mogroside V-predominant fermentation-derived monk fruit sweetener ingredient. The ingredient was determined to be non-mutagenic in the bacterial reverse mutation test in all five tester strains, with and without metabolic activation, up to the maximum recommended concentration of 5000 µg/plate, and was neither clastogenic nor aneugenic in cultured human peripheral blood lymphocytes up to 5000 µg/mL in all three treatment regimens. The 90-day study identified no adverse effects at any dose, giving a NOAEL of 2000 mg/kg body weight/day, the highest dose tested.

### 4.1. Interpretation of systemic findings

Daily oral gavage administration of the preparation at up to 2000 mg/kg body weight/day for 90 days produced no adverse effects in Sprague-Dawley rats. The study showed no mortality and no test article-related effects across key safety domains, including clinical signs, ophthalmology, neurobehavioural endpoints (functional observational battery and motor activity), body weight and body-weight gain, food consumption, oestrous cyclicity, and sperm parameters. Clinical pathology (haematology, coagulation, clinical chemistry), thyroid hormones, and urinalysis showed no toxicologically meaningful, treatment-related change, and there were no test article-related gross or microscopic findings in any tissue examined.

The only finding attributed to treatment was a modest increase in liver weight, observed in males at 2000 mg/kg/day and in females at 1000 and 2000 mg/kg/day. This finding was judged non-adverse because it occurred without any corroborating change in hepatocellular or hepatobiliary clinical chemistry markers (ALT, ALP, GGT, bilirubin) and without any hepatic gross or microscopic correlate. Equivalent findings, in each case without a histopathological correlate and in each case judged to be of no toxicological significance, have been reported for other monk fruit preparations: liver-weight changes in a monk fruit extract containing 52% mogroside V, which EFSA accepted as an adaptive, non-adverse response (EFSA, 2019); dose-dependent liver-weight increases in a fermentation-derived siamenoside I ingredient (Mihalchik et al., 2026); increased relative liver weight in male rats at the highest dietary concentration over 13 weeks (Jin et al., 2007); and increased absolute and relative liver weights in high-dose rats over 28 days (Marone et al., 2008). Because the study design did not include a recovery or withdrawal period, the reversibility of this change was not assessed.

The remaining statistically significant differences, scattered haematology (mean platelet volume, white cell, lymphocyte, and neutrophil counts; APTT), clinical chemistry (total bilirubin), thyroid (T4 in females), grip strength, foot splay, and interval body weight gain and food consumption, were small in magnitude, inconsistent across dose, time, and sex, unaccompanied by correlative functional or morphological changes, and within the laboratory historical control ranges. Such isolated differences are typical of the biological variability inherent in guideline subchronic studies and were considered incidental. Notably, the gavage design avoided the reductions in body weight gain seen across dose groups in dietary mogroside studies, where replacement of a portion of the diet with a non-caloric ingredient lowers caloric density (Marone et al., 2008; Mihalchik et al., 2026); the absence of any body-weight effect here indicates that those dietary findings reflect caloric dilution rather than toxicity.

### 4.2. Relevance to the male reproductive concern

Although monk fruit extracts and juice concentrates are widely used in Asia and are Generally Recognized as Safe (GRAS) in the United States, their regulatory position in the European Union is still pending. The European Food Safety Authority (EFSA) reviewed monk fruit extract as a food additive in 2019 and concluded that the available database was insufficient to establish its safety; in particular, the Panel could not exclude a potential concern related to decreases in testis weight, accompanied by histopathological findings, reported in a 90-day dietary study of a monk fruit extract containing 52% mogroside V (EFSA, 2019). A specific aqueous monk fruit extract has since been authorized as a novel food under Commission Implementing Regulation (EU) 2024/2345, but highly purified mogroside preparations remain unauthorized, reflecting persisting gaps in the toxicological database and the absence of industry-led applications (Kaim et al., 2025). Resolving those gaps requires additional guideline-compliant toxicological data on well-characterized, high-purity mogroside ingredients, with specific attention to male reproductive endpoints.

The present study, conducted with a mogroside V-predominant ingredient at high gavage doses, provides directly relevant reassurance: there were no test item-related effects on absolute or relative testis weight, on sperm motility, morphology, or counts, or on the qualitative staging of spermatogenesis, and no microscopic findings in any reproductive tissue. These results do not support a mogroside V-related effect on male reproductive organs under the conditions tested. Although the 90-day dosing period spans the full duration of spermatogenesis and subsequent epididymal transit in the rat, complex reproductive and developmental endpoints, including functional fertility, mating behaviour, and multigenerational outcomes, cannot be evaluated in a standard systemic toxicity study; such dedicated studies were outside the scope of the present work, and the findings reported here provide no indication of a male reproductive hazard.

### 4.3. Comparison with prior mogroside studies and estimated human exposure

The metabolism and pharmacokinetics of mogrosides have been examined in several *in vitro* and *in vivo* studies, which together describe a consistent disposition pathway. Mogroside V resists digestion in the upper gastrointestinal tract and is deglycosylated by intestinal microflora to the aglycone mogrol and its mono- and diglucosides, which are excreted predominantly in the feces; only trace quantities of mogrol and its conjugates reach the portal circulation (Murata et al., 2010). The mogrol that is absorbed undergoes metabolism and is eliminated primarily in the bile, with little excreted in the urine. Structurally distinct mogrosides, including mogroside V and siamenoside I, produce the same aglycone, mogrol, when metabolized irrespective of the number or linkage of glucose units (Bhusari et al., 2021; Xu et al., 2015), and single-dose pharmacokinetic data for siamenoside I confirm pre-absorptive deglycosylation to mogrol, first-pass oxidation, and rapid faecal elimination without tissue accumulation (Roberts et al., 2026).

Although the present ingredient is produced by fermentation rather than by extraction of monk fruit, the mogrosides it contains are structurally identical to those occurring naturally in *S. grosvenorii*, and its purity (at least 95% total mogrosides) reduces the co-extracted plant constituents that confound the interpretation of findings for conventional extracts. Its toxicological behavior would therefore be expected to be governed by the mogrosides and their common aglycone rather than by the production route, and the available data are consistent with that expectation: the ingredient evaluated here and the siamenoside I-rich ingredient of Mihalchik et al. (2026) differ both in the predominant mogroside and in fermentation host (*E. coli* versus *Y. lipolytica*), yet gave concordant outcomes across genotoxicity, clinical pathology, and histopathology.

In neither study was an effect level identified, both NOAELs having been set by the highest dose that could practically be administered.

The NOAEL of 2000 mg/kg body weight/day is lower in absolute terms than the values reported for other monk fruit and mogroside preparations, which range from approximately 2500 to 7500 mg/kg body weight/day (Supplementary Table S1d). In all mogroside-related studies, the NOAEL was the highest dose or dietary concentration tested, so none identifies an effect level for mogrosides. With the exception of the fermentation-derived ingredient of Mihalchik et al. (2026), the earlier studies used extracts or concentrates in which mogrosides accounted for only 62–86% of the administered material; expressed on a total-mogroside basis, the published NOAELs fall to approximately 2500–4600 mg/kg body weight/day against 1900 mg/kg body weight/day within the current study which on a mogroside V basis corresponds to 1410 mg/kg body weight/day.

Anticipated human exposure lies far below all of these levels. Because the preparation is approximately 250 times as sweet as sucrose, only small quantities are required to achieve the intended sweetness in the proposed food categories. Estimated daily intakes were derived using the sweetener substitution method of Renwick (2008), in which reported intakes of currently authorized high-intensity sweeteners, expressed as sucrose equivalents, are assumed to be replaced in their entirety by the new sweetener and adjusted for its relative sweetness, an approach that is conservative in assuming complete substitution and brand loyalty, while remaining grounded in measured post-market intakes. On this basis, estimated intakes range from 1.02 mg/kg body weight/day in average adult consumers to 3.96 mg/kg body weight/day in high-consumer children, the highest of four population groups considered; average and high-consumer estimates for diabetic adults and diabetic children fall within this range. The highest estimate is approximately 500-fold below the NOAEL established here, and approximately five-fold below the 20 mg/kg body weight/day that the NOAEL would support after application of the default 100-fold uncertainty factor.

## 5. Conclusions

Under the conditions of this GLP, OECD 408-compliant study, oral gavage administration of a high-purity, mogroside V-predominant preparation to Sprague-Dawley rats for 90 consecutive days was not associated with any treatment-related adverse effect, including male reproductive endpoints. The same preparation was non-mutagenic in the bacterial reverse mutation test across five tester strains and neither clastogenic nor aneugenic in the in vitro human lymphocyte micronucleus test, in each case up to the maximum recommended concentration and both with and without metabolic activation. The no-observed-adverse-effect level (NOAEL) was 2000 mg/kg body weight/day, the highest dose tested. Together with the complementary data on a siamenoside I-rich fermentation-derived ingredient (Mihalchik et al., 2026), these results support the safety of high-purity mogroside preparations for use as natural, non-nutritive sweetening ingredients. As sugar-reduction efforts increase demand for natural high-potency sweeteners, fermentation provides a scalable, geographically resilient route to monk fruit sweeteners that are independent of monk fruit agriculture concentrated in a single region, while delivering a defined, high-purity ingredient. Because the fermentation-produced mogrosides are structurally identical to those of *S. grosvenorii*, are metabolized by the same route, undergoing pre-absorptive deglycosylation by the intestinal microflora to the common aglycone mogrol, and the ingredient contains substantially less co-extracted plant material than conventional extracts, the safety demonstrated here both draws on, and adds a more comprehensively characterized dataset to, the existing knowledge base for monk fruit sweeteners.

## CRediT authorship contribution statement

Christine Nicole S. Santos: Funding acquisition, Validation, Visualization, Resources, Writing – original draft, Writing – review & editing. Sulaiman S M: Investigation, Formal analysis, Data curation, Validation. Sanjaykumar M. Paneliya: Investigation, Formal analysis, Data curation, Validation. Martyna Pawluk – Conceptualization, Validation, Writing – review & editing. Srikanth A – Project administration, Supervision, Resources. Venkata Sai Prakash Chaturvedula – Conceptualization, Validation, Writing – review & editing.

## Data availability

Data will be made available on request.

### Acknowledgements

The authors thank the staff of Adgyl Lifesciences Private Limited for conduct of the in-life, clinical pathology, and pathology phases of the study.

## Declaration of generative AI and AI-assisted technologies in the writing process

During the preparation of this work, the authors used Claude (Anthropic; model version Opus 5) in order to extract and tabulate clinical pathology and genotoxicity data from the underlying study reports into the manuscript and supplementary tables, to cross-check tabulated values against those reports, and to assist with editing of the manuscript text. After using this tool, the authors reviewed and edited the content as needed, verified all reported data against the original study reports, and take full responsibility for the content of the publication.

