## Supplementary Tables for "Genotoxicity and 90-day oral toxicity of a monk fruit (*Siraitia grosvenorii*) mogroside preparation (≥95% mogrosides) produced by microbial fermentation"

**Table S1a.** Complete haematology and coagulation panel at termination of the 90-day oral toxicity study (Day 91). Dose levels are given in mg/kg body weight/day.

| Parameter | Unit | Males (n = 10 per group) | | | | Females (n = 10 per group) | | | |
|---|---|---|---|---|---|---|---|---|---|
| | | 0 | 500 | 1000 | 2000 | 0 | 500 | 1000 | 2000 |
| Red blood cell count (RBC) | $10^{12}$/L | 9.08 ± 0.31 | 9.02 ± 0.29 | 8.77 ± 0.32* | 9.11 ± 0.23 | 8.19 ± 0.26 | 8.27 ± 0.38 | 8.25 ± 0.28 | 8.28 ± 0.22 |
| Haemoglobin (HGB) | g/L | 168 ± 5 | 168 ± 6 | 168 ± 8 | 172 ± 4 | 157 ± 5 | 160 ± 4 | 161 ± 6 | 160 ± 4 |
| Haematocrit (HCT) | L/L | 0.522 ± 0.012 | 0.527 ± 0.019 | 0.518 ± 0.021 | 0.536 ± 0.014 | 0.491 ± 0.015 | 0.493 ± 0.010 | 0.502 ± 0.017 | 0.493 ± 0.011 |
| Mean cell volume (MCV) | fL | 57.5 ± 1.6 | 58.5 ± 2.3 | 59.1 ± 1.1 | 58.9 ± 1.1 | 60.0 ± 1.7 | 59.6 ± 2.2 | 60.9 ± 1.6 | 59.5 ± 1.3 |
| Mean cell haemoglobin (MCH) | pg | 18.5 ± 0.7 | 18.6 ± 0.7 | 19.1 ± 0.6 | 18.9 ± 0.4 | 19.2 ± 0.4 | 19.4 ± 0.8 | 19.5 ± 0.6 | 19.3 ± 0.5 |
| Mean cell haemoglobin concentration (MCHC) | g/L | 322 ± 7 | 318 ± 4 | 323 ± 6 | 321 ± 7 | 319 ± 3 | 325 ± 5* | 321 ± 5 | 324 ± 5 |
| Reticulocytes, absolute | $10^{12}$/L | 0.206 ± 0.033 | 0.213 ± 0.029 | 0.200 ± 0.033 | 0.201 ± 0.016 | 0.225 ± 0.036 | 0.200 ± 0.024 | 0.230 ± 0.053 | 0.200 ± 0.034 |
| Red cell distribution width (RDW) | % | 12.3 ± 0.4 | 12.7 ± 0.4 | 12.7 ± 0.5 | 12.5 ± 0.3 | 11.5 ± 0.3 | 11.5 ± 0.3 | 11.9 ± 0.3* | 11.8 ± 0.3 |
| Haemoglobin distribution width (HDW) | g/L | 22.5 ± 1.1 | 22.8 ± 1.5 | 23.2 ± 1.6 | 22.7 ± 1.4 | 20.6 ± 0.9 | 21.2 ± 1.4 | 21.3 ± 1.2 | 20.8 ± 1.1 |
| Hyperchromic erythrocytes | % | 0.7 ± 0.4 | 0.6 ± 0.5 | 0.7 ± 0.6 | 0.4 ± 0.2 | 0.4 ± 0.2 | 0.7 ± 0.5 | 0.2 ± 0.1 | 0.3 ± 0.3 |
| Hypochromic erythrocytes | % | 0.2 ± 0.1 | 0.2 ± 0.1 | 0.2 ± 0.1 | 0.2 ± 0.1 | 0.1 ± 0.0 | 0.1 ± 0.0 | 0.2 ± 0.1* | 0.1 ± 0.0 |
| Macrocytic erythrocytes | % | 0.0 ± 0.0 | 0.0 ± 0.0 | 0.0 ± 0.0 | 0.0 ± 0.0 | 0.0 ± 0.0 | 0.0 ± 0.0 | 0.0 ± 0.0 | 0.0 ± 0.0 |
| Microcytic erythrocytes[a] | % | 0.0 ± 0.0 | 0.0 ± 0.0 | 0.0 ± 0.0 | 0.0 ± 0.0 | 0.0 ± 0.0 | 0.0 ± 0.0 | 0.0 ± 0.0 | 0.0 ± 0.0 |
| Red cell fragments | $10^{12}$/L | 0.14 ± 0.02 | 0.15 ± 0.02 | 0.17 ± 0.04* | 0.13 ± 0.02 | 0.14 ± 0.02 | 0.15 ± 0.02 | 0.14 ± 0.02 | 0.15 ± 0.02 |
| Red cell ghosts[a] | $10^{12}$/L | 0.00 ± 0.00 | 0.00 ± 0.00 | 0.00 ± 0.00 | 0.00 ± 0.00 | 0.00 ± 0.00 | 0.00 ± 0.00 | 0.00 ± 0.00 | 0.00 ± 0.00 |
| Platelet count (PLT) | $10^9$/L | 1283 ± 110 | 1295 ± 63 | 1483 ± 304 | 1190 ± 131 | 1289 ± 140 | 1363 ± 124 | 1315 ± 121 | 1312 ± 157 |
| Mean platelet volume (MPV) | fL | 9.6 ± 0.4 | 10.3 ± 0.5* | 10.5 ± 0.3* | 10.8 ± 0.4* | 10.1 ± 0.4 | 10.4 ± 0.3 | 10.6 ± 0.3* | 10.5 ± 0.3* |
| White blood cell count (WBC) | $10^9$/L | 10.09 ± 1.48 | 10.20 ± 1.51 | 9.70 ± 1.82 | 8.15 ± 1.89* | 7.06 ± 1.08 | 5.69 ± 1.02* | 5.98 ± 1.07 | 7.62 ± 1.61 |
| Neutrophils, absolute | $10^9$/L | 1.79 ± 0.38 | 2.14 ± 0.96 | 1.74 ± 0.51 | 1.61 ± 0.50 | 1.02 ± 0.18 | 0.90 ± 0.23 | 1.32 ± 1.41 | 1.30 ± 0.52 |
| Lymphocytes, absolute | $10^9$/L | 7.73 ± 1.45 | 7.50 ± 1.00 | 7.45 ± 1.42 | 6.09 ± 1.58* | 5.64 ± 0.89 | 4.44 ± 1.06 | 4.26 ± 1.42* | 5.91 ± 1.33 |
| Monocytes, absolute | $10^9$/L | 0.29 ± 0.08 | 0.31 ± 0.06 | 0.25 ± 0.10 | 0.21 ± 0.07 | 0.22 ± 0.10 | 0.17 ± 0.03 | 0.25 ± 0.25 | 0.20 ± 0.13 |
| Basophils, absolute | $10^9$/L | 0.03 ± 0.01 | 0.03 ± 0.01 | 0.02 ± 0.01 | 0.02 ± 0.01* | 0.02 ± 0.01 | 0.02 ± 0.01 | 0.01 ± 0.00* | 0.02 ± 0.01 |
| Eosinophils, absolute | $10^9$/L | 0.18 ± 0.08 | 0.15 ± 0.03 | 0.16 ± 0.04 | 0.16 ± 0.08 | 0.11 ± 0.05 | 0.12 ± 0.04 | 0.08 ± 0.05 | 0.14 ± 0.09 |
| Prothrombin time (PT) | s | 19.3 ± 1.0 | 19.1 ± 0.9 | 19.5 ± 2.1 | 21.5 ± 3.2 | 16.8 ± 0.6 | 16.9 ± 0.8 | 16.9 ± 0.7 | 16.6 ± 0.9 |
| Activated partial thromboplastin time (APTT) | s | 17.4 ± 2.5 | 17.5 ± 2.6 | 17.4 ± 3.5 | 21.2 ± 2.3* | 12.9 ± 1.8 | 11.7 ± 2.4 | 11.0 ± 3.7 | 13.4 ± 1.5 |

*Blood was collected at scheduled termination (Day 91) from all animals after an overnight fast; haematology was performed on an ADVIA 2120i analyser and coagulation parameters on citrated plasma using a Ceveron Alpha analyser. Values are group mean ± SD. Group means were compared with the concurrent vehicle control by one-way analysis of variance followed by Dunnett's two-sided test, applied to untransformed, log-transformed or rank-transformed data according to the variance structure of each parameter; * denotes $p < 0.05$ versus control. [a] Statistical analysis was not appropriate for this parameter because all group values were zero. None of the differences marked was considered test item-related. Each was of small magnitude, lacked a consistent dose-response relationship or was inconsistent between the sexes, was unaccompanied by any correlative functional, organ-weight or morphological finding, and fell within the laboratory historical control range for the parameter concerned.*

**Table S1b.** Complete clinical chemistry panel at termination of the 90-day oral toxicity study (Day 91). Dose levels are given in mg/kg body weight/day.

| Parameter | Unit | Males (n = 10 per group) | | | | Females (n = 10 per group) | | | |
|---|---|---|---|---|---|---|---|---|---|
| | | 0 | 500 | 1000 | 2000 | 0 | 500 | 1000 | 2000 |
| Glucose | mmol/L | 6.38 ± 1.01 | 6.35 ± 0.81 | 6.13 ± 0.44 | 6.66 ± 0.71 | 6.92 ± 1.18 | 6.56 ± 0.64 | 6.50 ± 0.56 | 6.98 ± 0.84 |
| Blood urea nitrogen (BUN) | mmol/L | 4.72 ± 0.56 | 4.66 ± 0.45 | 4.60 ± 0.58 | 4.52 ± 0.30 | 5.38 ± 0.83 | 5.47 ± 0.77 | 5.85 ± 0.76 | 5.33 ± 0.75 |
| Creatinine | µmol/L | 40 ± 2 | 41 ± 4 | 42 ± 4 | 42 ± 4 | 45 ± 5 | 48 ± 3 | 50 ± 3* | 46 ± 4 |
| Aspartate aminotransferase (AST) | U/L | 94 ± 12 | 101 ± 39 | 94 ± 8 | 88 ± 11 | 100 ± 43 | 92 ± 19 | 142 ± 95 | 118 ± 66 |
| Alanine aminotransferase (ALT) | U/L | 32 ± 5 | 46 ± 38 | 31 ± 5 | 35 ± 11 | 30 ± 21 | 28 ± 14 | 45 ± 46 | 39 ± 43 |
| Gamma-glutamyl transferase (GGT)[a] | U/L | Below LLOQ | Below LLOQ | Below LLOQ | Below LLOQ | Below LLOQ | Below LLOQ | Below LLOQ | Below LLOQ |
| Alkaline phosphatase (ALP) | U/L | 85 ± 19 | 91 ± 15 | 78 ± 15 | 81 ± 16 | 57 ± 19 | 39 ± 5 | 36 ± 9* | 43 ± 17 |
| Creatine kinase (CK) | U/L | 147 ± 42 | 175 ± 37 | 208 ± 99 | 178 ± 63 | 121 ± 30 | 165 ± 129 | 153 ± 56 | 145 ± 53 |
| Total bilirubin | µmol/L | 2.31 ± 0.36 | 2.05 ± 0.62 | 1.92 ± 0.27 | 1.93 ± 0.45 | 2.96 ± 0.77 | 2.51 ± 0.27 | 2.38 ± 0.40 | 2.04 ± 0.37* |
| Total cholesterol | mmol/L | 1.64 ± 0.42 | 1.75 ± 0.28 | 1.52 ± 0.23 | 1.61 ± 0.29 | 2.32 ± 0.47 | 2.44 ± 0.40 | 2.33 ± 0.69 | 2.26 ± 0.47 |
| HDL cholesterol | mmol/L | 0.40 ± 0.06 | 0.43 ± 0.04 | 0.38 ± 0.06 | 0.41 ± 0.05 | 0.54 ± 0.06 | 0.57 ± 0.06 | 0.55 ± 0.08 | 0.55 ± 0.06 |
| LDL cholesterol | mmol/L | 0.26 ± 0.08 | 0.29 ± 0.04 | 0.28 ± 0.06 | 0.28 ± 0.06 | 0.29 ± 0.08 | 0.31 ± 0.07 | 0.31 ± 0.11 | 0.33 ± 0.13 |
| Triglycerides | mmol/L | 0.69 ± 0.51 | 0.68 ± 0.32 | 0.49 ± 0.20 | 0.69 ± 0.29 | 0.40 ± 0.14 | 0.28 ± 0.09 | 0.39 ± 0.26 | 0.34 ± 0.20 |
| Total protein | g/L | 71.0 ± 3.1 | 72.6 ± 2.3 | 70.0 ± 2.5 | 72.9 ± 1.7 | 79.3 ± 4.5 | 80.6 ± 1.9 | 81.6 ± 4.1 | 79.3 ± 4.2 |
| Albumin | g/L | 35.5 ± 1.2 | 36.7 ± 1.1* | 35.6 ± 0.9 | 36.8 ± 0.7* | 43.0 ± 3.1 | 43.0 ± 1.3 | 43.5 ± 2.7 | 41.1 ± 2.7 |
| Globulin (calculated) | g/L | 35.6 ± 2.1 | 35.9 ± 1.8 | 34.4 ± 2.0 | 36.1 ± 1.7 | 36.3 ± 2.0 | 37.6 ± 1.1 | 38.1 ± 2.0 | 38.2 ± 2.6 |
| Albumin:globulin ratio | ratio | 1.00 ± 0.04 | 1.02 ± 0.06 | 1.04 ± 0.06 | 1.02 ± 0.05 | 1.18 ± 0.08 | 1.14 ± 0.04 | 1.14 ± 0.07 | 1.08 ± 0.09* |
| Inorganic phosphorus | mmol/L | 1.94 ± 0.14 | 1.77 ± 0.14 | 1.80 ± 0.15 | 1.86 ± 0.18 | 1.60 ± 0.25 | 1.49 ± 0.19 | 1.36 ± 0.24 | 1.61 ± 0.19 |
| Calcium | mmol/L | 2.56 ± 0.05 | 2.57 ± 0.05 | 2.56 ± 0.06 | 2.58 ± 0.05 | 2.69 ± 0.10 | 2.70 ± 0.06 | 2.68 ± 0.10 | 2.70 ± 0.09 |
| Sodium | mEq/L | 143.5 ± 1.2 | 142.9 ± 1.1 | 142.2 ± 1.1 | 142.8 ± 1.0 | 141.4 ± 1.0 | 141.0 ± 0.9 | 140.9 ± 1.2 | 140.9 ± 1.2 |
| Potassium | mEq/L | 3.59 ± 0.25 | 3.61 ± 0.21 | 3.54 ± 0.21 | 3.63 ± 0.19 | 3.33 ± 0.18 | 3.25 ± 0.19 | 3.13 ± 0.15 | 3.44 ± 0.27 |
| Chloride | mEq/L | 104.0 ± 1.3 | 103.0 ± 0.8 | 102.7 ± 0.8* | 103.5 ± 1.0 | 103.1 ± 1.3 | 102.5 ± 0.9 | 101.9 ± 1.6 | 102.3 ± 1.6 |

*Blood was collected at scheduled termination (Day 91) from all animals after an overnight fast; clinical chemistry was performed on a Beckman Coulter AU480 analyser. Values are group mean ± SD. Group means were compared with the concurrent vehicle control by one-way analysis of variance followed by Dunnett's two-sided test, applied to untransformed, log-transformed or rank-transformed data according to the variance structure of each parameter; * denotes $p < 0.05$ versus control. [a] All individual gamma-glutamyl transferase values in every group, including controls, were below the lower limit of quantification of 5 U/L; no group mean could therefore be derived and the parameter was not analysed statistically. None of the differences marked was considered test item-related. Each was of small magnitude, lacked a consistent dose-response relationship or was inconsistent between the sexes, was unaccompanied by any correlative functional, organ-weight or morphological finding, and fell within the laboratory historical control range for the parameter concerned.*

**Table S1c.** Quantitative urinalysis parameters at termination of the 90-day oral toxicity study (Day 91). Dose levels are given in mg/kg body weight/day.

| Parameter | Unit | Males (n = 10 per group) | | | | Females (n = 10 per group) | | | |
|---|---|---|---|---|---|---|---|---|---|
| | | 0 | 500 | 1000 | 2000 | 0 | 500 | 1000 | 2000 |
| Urine volume | mL | 38.9 ± 8.9 | 39.5 ± 9.2 | 23.8 ± 7.3* | 26.8 ± 8.3* | 20.4 ± 8.4 | 18.5 ± 5.0 | 11.5 ± 2.6* | 16.9 ± 5.6 |
| Urine pH | - | 8.6 ± 0.2 | 8.5 ± 0.0 | 8.5 ± 0.2 | 8.0 ± 0.4* | 8.5 ± 0.0 | 8.5 ± 0.0 | 8.3 ± 0.3 | 8.3 ± 0.4* |
| Specific gravity | - | 1.010 ± 0.004 | 1.012 ± 0.004 | 1.015 ± 0.005 | 1.016 ± 0.004* | 1.013 ± 0.005 | 1.016 ± 0.003 | 1.023 ± 0.004* | 1.018 ± 0.006 |

*Urine was collected overnight (Day 90-91) from all animals in metabolism cages before the terminal blood collection. Values are group mean ± SD. Group means were compared with the concurrent vehicle control by one-way analysis of variance followed by Dunnett's two-sided test, applied to untransformed, log-transformed or rank-transformed data according to the variance structure of each parameter; * denotes $p < 0.05$ versus control. Qualitative and semi-quantitative parameters (colour, clarity, nitrite, protein, glucose, ketones, urobilinogen, bilirubin, erythrocytes, leukocytes and microscopic sediment) were recorded as incidences rather than as group means and showed no test item-related pattern; they are not reproduced here. The lower urine volume and correspondingly higher specific gravity at 1000 and 2000 mg/kg body weight/day were unaccompanied by any change in blood urea nitrogen, creatinine or electrolytes, or by any kidney organ-weight or histopathological correlate, and are consistent with a difference in urine concentrating status rather than with renal injury.*

**Table S2.** Comparison with published repeated-dose toxicity studies of monk fruit and mogroside preparations, expressed both as administered and on a mogroside-equivalent basis.

| Study | Test article (reported composition) | Species, route, duration | NOAEL as administered | NOAEL as total mogrosides | NOAEL as predominant mogroside |
|---|---|---|---|---|---|
| Present study | Fermentation-derived; ≥95% total mogrosides, 70.3% mogroside V | Rat, gavage, 90 days | 2000 (M and F) | 1900 | 1410 (mogroside V) |
| Mihalchik et al. (2026) | Fermentation-derived; ≥91.9% total mogrosides, 75.8% siamenoside I | Rat, diet, 90 days | 3309 (M) 3362 (F); TWA | 3040, 3090 | 2510, 2550 (siamenoside I) |
| Marone et al. (2008) | Luo Han fruit concentrate; approx. 62% total mogrosides, approx. 39% mogroside V | Rat, diet, 28 days | 7070 (M) 7480 (F) | 4380, 4640 | 2760, 2920 (mogroside V) |
| Jin et al. (2007) | S. grosvenorii extract; composition not reported | Rat, diet, 13 weeks | 2520 (M) 3200 (F) | Not calculable | Not calculable |
| Qin et al. (2006) | Mogroside extract; 81.5-85.7% total mogrosides, >30% mogroside V | Dog, gavage, 90 days | 3000 (M and F) | approx. 2510 | >900 (mogroside V) |

*All values in mg/kg body weight/day. In every study the NOAEL was the highest dose or dietary concentration tested, so none of these values identifies an effect level. Mogroside-equivalent doses are calculated by applying the test article composition reported in each paper to the NOAEL as administered, and are approximate because the reported compositions are themselves ranges or lower bounds. TWA, time-weighted average; M, males; F, females.*
